# Early life psychosocial stress increases binge-like ethanol consumption and CSF1R inhibition prevents stress-induced alterations in microglia and brain macrophage population density

**DOI:** 10.1101/2024.07.27.605403

**Authors:** Stephen C. Gironda, Samuel W. Centanni, Jeffrey L. Weiner

**Affiliations:** Department of Translational Neuroscience, Wake Forest University School of Medicine, Winston Salem, North Carolina, 27101

**Keywords:** Early Life Stress, Alcohol, Microglia, Light-sheet Fluorescence Microscopy, CSF1R inhibition, Macrophage

## Abstract

Early life stress (ELS) has lasting consequences on microglia and brain macrophage function. During ELS, microglia and brain macrophages alter their engagement with synapses leading to changes in neuronal excitability. Further, ELS can induce innate immune memory formation in microglia and brain macrophages resulting in altered responsivity to future environmental stimuli. These alterations can result in lasting adaptations in circuit function and may mediate the relationship between ELS and the risk to develop alcohol use disorder (AUD). Whether microglia and brain macrophages truly mediate this relationship remains elusive. Here, we report: 1) an ELS model, psychosocial stress (PSS), increases binge-like ethanol consumption in early adulthood. 2) Repeated binge-like ethanol consumption increases microglia and brain macrophage population densities across the brain. 3) PSS may elicit innate immune memory formation in microglia and brain macrophages leading to altered population densities following repeated binge-like ethanol consumption. 4) Microglia and brain macrophage inhibition trended towards preventing PSS-evoked changes in binge-like ethanol consumption and normalized microglia and brain macrophage population densities. Therefore, our study suggests that acutely inhibiting microglia and brain macrophage function during periods of early life PSS may prevent innate immune memory formation and assist in reducing the risk to develop AUD.

**Highlights:**

1. An early life psychosocial stress (PSS) exposure increases ethanol consumption
2. Microglial inhibition during PSS trends towards reducing ethanol consumption
3. Binge ethanol consumption increases microglial count and alters cell proximity
4. Early life PSS alters microglial responsivity to binge ethanol consumption
5. Microglial inhibition may prevent microglial innate immune memory formation

## 1. Introduction

Over 63% of the US population has experienced early life stress (ELS) (Swedo et al., 2023). ELS encompasses a variety of early life experiences including sexual, emotional, and physical abuse (Wade et al., 2022). Consequences of ELS include rapid brain development, dysregulated executive function, and issues with emotion-regulation and reward appraisal (Herzberg & Gunnar, 2020). Further, ELS is a risk factor for a host of neuropsychiatric disorders including alcohol use disorder (AUD) (Kirsch & Lippard, 2022). One form of ELS, bullying victimization, is known to have negative effects on the brain and behavior (Bhatia et al., 2023). Bullying is characterized as aggressive behavior that is repetitive and intentional in which a power differential exists between the bully and the victim (Armitage, 2021). An FMRI study investigating the effect of adolescent bullying on brain activity and suicide risk demonstrated that social threat cues related to bullying increased activation of numerous brain regions including the insula, anterior cingulate cortex, and amygdala (Yang et al., 2023). Studies also suggest bullying victimization can increase the risk for binge drinking (Davis et al., 2018) and the development of AUD (Hallit et al., 2020).

Alcohol use remains one of the highest-ranked causes of death worldwide. Excessive and maladaptive alcohol use, clinically defined as AUD, affects nearly 29.6 million people in the United States (SAMHSA, 2023). Excessive alcohol use costs the US roughly $249 billion dollars a year. A vast majority of these costs ($191.1 billion) are accrued through a particularly dangerous behavior associated with AUD: binge drinking (Sacks et al., 2015). Identifying and characterizing the effects of ELS on the brain can provide clinicians with innovative approaches to prevent the onset of binge drinking, and more broadly AUD.

ELS can induce alterations in reward circuit function that may lead to increased motivation for, and consumption of alcohol (Koob & Volkow, 2010, Hanson et al., 2021). Although studies have documented these ELS-mediated circuit adaptations, less is known about how these adaptations occur. Several preclinical models are used to explore the effects of ELS on alcohol consumption and neurobiology. These include maternal separation (Bertagna et al., 2021), adolescent social isolation (Chappell et al., 2013; Whitaker et al., 2013; Skelly et al., 2015; Butler et al., 2016), limited bedding and nesting (Okhuarobo et al., 2020), adverse social experiences (Surakka et al., 2021), and social defeat (Rodriguez-Arias et al., 2016).

While results from these studies show a variety of consequences on behavior and neurobiology, few have been implemented between postnatal days (P) 14-21. This period of postnatal development is critical for the cortex (Andolina et al., 2011), the immune system (Danese & Lewis, 2017), and synapse maturation (Paolicelli et al., 2011). Recent work has investigated this developmental window by employing a translationally relevant model of bullying victimization, early life psychosocial stress (PSS). Findings from this work suggest that PSS can induce drug-seeking behaviors in mice; and the combination of PSS and cocaine treatment increases microglia count and fluorescence intensity (Lo Iacono et al., 2018). But what are microglia?

Microglia and brain macrophages are vital to healthy brain development and homeostasis (Paolicelli et al., 2022). During development, microglia and brain macrophages regulate synapse maturation and elimination to ensure proper neuron and circuit function (Paolicelli & Ferretti, 2017; Bolton et al., 2022; Surala et al., 2024; Zhao et al., 2024). ELS can alter these developmental processes and change microglia and brain macrophage function longitudinally (Catale et al., 2020; Bolton et al., 2022). This process, referred to as innate immune memory formation or priming, can result in altered microglial and brain macrophage responsivity to future environmental stimuli (Frank et al., 2014; Haley et al., 2019; Neher & Cunningham, 2019, Reemst et al., 2022). Recent work speculates that microglial priming could play a role in the development of AUD (Melbourne et al., 2021). As a result, inhibiting microglia and brain macrophage innate immune memory formation during ELS may provide a promising approach to prevent the immediate consequences of ELS on synapse maturation and the long-term consequences of ELS on circuit function and the risk to develop AUD.

In this study, we exposed mice to PSS from P14-21. In early adulthood, mice performed the drinking in the dark (DID) paradigm to determine whether PSS could alter binge-like ethanol consumption. A subset of PSS-exposed mice received a colony stimulating factor 1 receptor (CSF1R) inhibitor, GW2580, during the stress exposure to determine whether microglia and brain macrophage inhibition could prevent alterations in binge-like ethanol consumption. We then characterized the effect of PSS, repeated binge-like ethanol consumption, and GW2580 treatment on microglia and brain macrophage population densities across twelve brain regions using light-sheet fluorescence microscopy (LSFM).

## 2. Methods and Materials

### 2.1 Animals

Ten-week-old Hsd:ICR CD-1 outbred mice (CD1) male (n=10) (Inotiv Co), and six-week-old male (n=4) and female (n=8) B6.129P2(Cg)-Cx3cr1tm1Litt/J (CX3CR1^GFP+^) mice were purchased from Jackson Laboratories (Strain #:005582). C- X3-C motif receptor 1 (CX3CR1) regulates cell motility in response to changes in environmental stimuli (Lee et al., 2018). Further, the CX3CR1^GFP+^ mouse line is highly specific for microglia and macrophages in the brain (Chakraborty et al., 2019). Thus, this mouse line is ideal for investigating brain-wide changes in microglia and brain macrophage population densities with techniques like LSFM (Müllenbroich et al., 2018). Food and water were provided ad libitum, and mice were housed on a 12:12 light:dark cycle with lights on at 07:00 AM. Behavioral paradigms were performed during lights off from 9:30 PM to 3:30 AM. All experimental procedures were approved by the Institutional Animal Care and Use Committee (IACUC) at Wake Forest University.

### 2.2 Orchiectomies

Testes were removed from adult CD1 male mice to deplete testosterone levels and reduce the likelihood that PSS-exposed pups would experience physical harm (Lo Iacono et al., 2017). Briefly, mice were anesthetized using vaporized isoflurane and placed on top of a heating pad. Mice were shaved using electric clippers along the abdominal cavity. Skin was then swabbed with 70% ethanol followed by sterile PBS. Surgical instruments including forceps and surgical scissors were sterilized prior to each surgery. A horizontal incision was then made an inch above the scrotum and forceps were used to free the testicular fat pad. Testes were then removed using surgical scissors and the remaining fat pad was returned to the scrotum. The incision was then closed using an intermittent suturing technique. Antibiotic ointment was applied to the incision location, and mice were returned to single-housed sterile cages upon recovery from anesthesia. CD1 mice were provided 30 days to recover from the surgery. Prior to the start of the PSS exposure, CD1 mice were tested for aggressive behaviors to confirm there was no threat of physical harm for the CX3CR1^GFP+^ pups.

### 2.3 Early Life Psychosocial Stress (PSS) Procedure

Each CX3CR1^GFP+^ male was mated with two CX3CR1^GFP+^ female mice at eight weeks of age. Once pregnancy was determined, female CX3CR1^GFP+^ breeders were moved to single-housing until the litter was born. Litters of CX3CR1^GFP+^ pups were randomly assigned to +/- PSS prior to P14. From P14-21, pups designated for PSS were briefly separated from the dam and litter; and placed in the home cage of a single-housed adult CD1 male mouse for 30 minutes each evening of the exposure. After each 30 minute PSS exposure, pups were returned to their dam and left undisturbed until the next evening. For each evening of the exposure, PSS-exposed pups were paired with a different adult CD1 mouse to control for individual differences in CD1 mouse behavior. Following PSS on P22, control and PSS-exposed pups were weaned and group-housed with pups of a similar age and from the same exposure condition until early adulthood (P60) (Lo Iacono et al., 2016, 2017, 2018).

### 2.4 GW2580 administration

A randomly assigned subset of CX3CR1 ^GFP+^ mice selected for PSS were administered a CSF1R inhibitor, GW2580. GW2580 is a selective antagonist of CSF1Rs which are found on microglia and myeloid lineage cells. Antagonism of these receptors inhibits microglial proliferation resulting in transient reductions in cell number acutely and ablation over longer administration windows (Neal et al., 2020). GW2580 (Selleck Chem, Catalog #: S8042) was suspended in a sterile saline buffer containing 0.5% hydroxypropylmethylcellulose and 0.1% Tween 80 (Martínez-Muriana et al., 2016). Control (n=21; 13 male and 8 female) and PSS-exposed (n=18; 9 male and 9 female) mice were administered a vehicle control, while another set of PSS-exposed mice (n=18; 10 male and 8 female) were administered GW2580 (75mg/kg). Intraperitoneal injections of vehicle and GW2580 were provided 30 minutes prior to each PSS exposure (Lo Iacono et al., 2018).

### 2.5 Drinking in the Dark (DID) procedure

At P60, mice were exposed to a modified DID procedure. Previous work suggests that this model of binge-like ethanol consumption promotes a clinically relevant point of intoxication in mice (Rhodes et al., 2005; Thiele & Navarro, 2014). Briefly, mice were single-housed in new cages without food or water two hours after lights off. Mice were provided one hour prior to the start of DID to acclimate to the single-housing. Once acclimated, sipper tubes containing 20% w/v ethanol were provided to the mice. On the first three nights of the procedure, mice were provided two hours of ad libitum ethanol access. On the fourth night, mice received a four hour extended access window. Following the four day procedure, mice were provided three days of abstinence. This procedure was repeated across four weeks. Each night of the exposure, mice were weighed, and the amount of ethanol consumed was measured as grams of ethanol (g) per kilogram (kg) of body weight (See Figure 1 for the experimental timeline).

**Figure 1.**
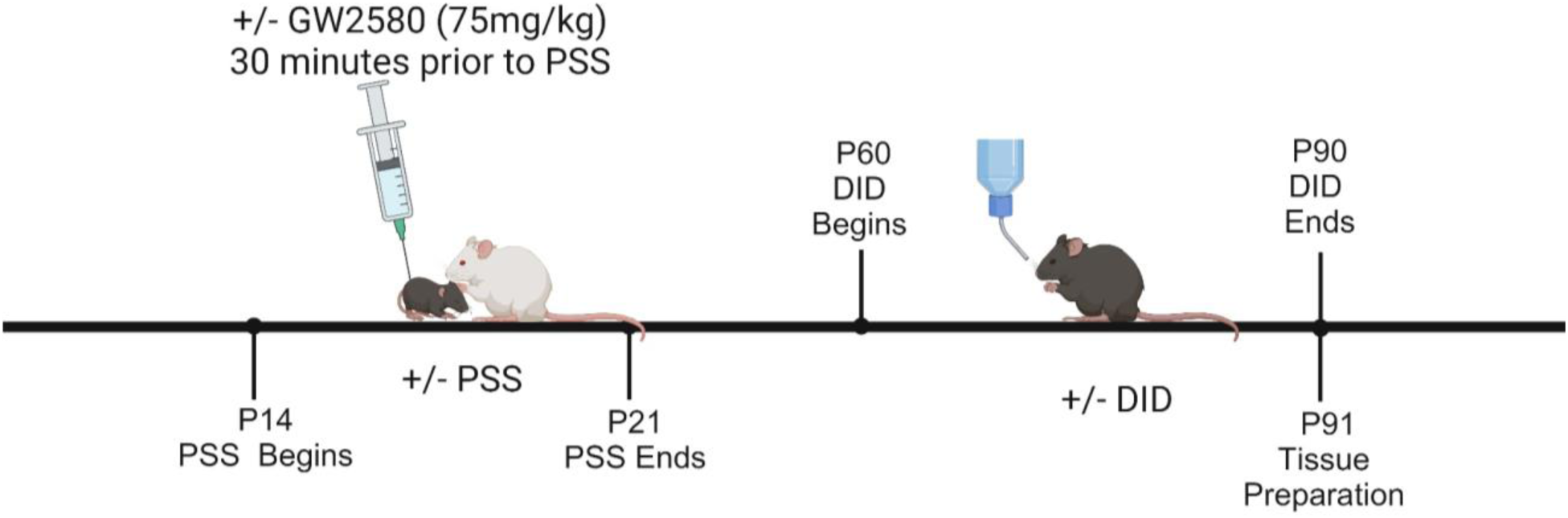
Experimental timeline: CX3CR1^GFP+^ pups were randomly assigned to either control or PSS at P14. Control (n=21) and PSS-exposed (n=18) mice were administered a vehicle, while a cohort of PSS-exposed (n=18) mice were administered GW2580 30 minutes prior to each PSS exposure. Following PSS at P22, control and PSS-exposed pups were weaned and group-housed until P60. At P60, control and PSS-exposed mice performed the DID procedure. 24 hours following the final DID exposure (P91), tissue was collected for LSFM.

### 2.6 Tissue Preparation for Light-sheet Fluorescence Microscopy (LSFM) Experiments

24 hours following completion of the DID procedure (P91), CX3CR1^GFP+^ mice were anesthetized using euthasol and prepared for transcardial perfusion. A vertical incision was made along the ventral mid-line of the mouse. The sternum and rib cage were removed to expose the heart. A 23-gauge butterfly needle was attached to a perfusion pump and was used to puncture the left ventricle of the heart. Ice cold dPBS + 10 mg/L heparin was flushed into the system at a flow rate of 8mLs/minute for three minutes. Following the dPBS flush, 4% PFA + 10mg/L heparin was perfused at 8mLs/minute for three minutes. Brains were extracted and stored in 4% PFA in PBS for 24 hours at 4° Celsius before preservation with a stabilization to harsh conditions via intramolecular epoxide linkages to prevent degradation (SHIELD) buffer kit (Park et al., 2018, Luschsinger et al., 2021).

Following preservation, a brain clearing solution containing sodium dodecyl sulfate (final concentration 300 mM), boric acid (final concentration 10mM), sodium sulfite (final concentration 100mM) and sodium hydroxide (titrated until pH 9) in Milli-q filtered water was prepared. Whole CX3CR1^GFP+^ brains were incubated in this solution at 37°C with gentle rocking for 20 days to ensure the tissue was transparent and to minimize light absorption and scattering. Once cleared, tissue was refraction indexed- matched using Easy Index (Life Canvas Technologies) to prevent optical distortion during image capture. Brains were then mounted on a sample holder with an Easy Index agarose gel solution (.45g of agarose/40mL of easy index) as stated per the manufacturer instructions (www.Lifecanvastech.com).

### 2.7 Light-sheet Fluorescence Microscopy (LSFM) Imaging

Following brain clearing, brains were mounted on a LifeCanvas SmartSPIM light-sheet microscope. Samples were illuminated with excitation wavelengths at 488nm to excite for CX3CR1^GFP+^ and 561nm as a blank control channel to account for any background or non-specific fluorescence. Laser power was adjusted for each brain and channel. Images were captured at 4 µm steps. Following image capture, raw files were migrated to a local server for destripping and stitching. Stitched images were stored as TIFFs and converted into Imaris files using Imaris File Converter 10.1.

### 2.8 Light-sheet Fluorescence Microscopy (LSFM) Analyses

Imaris files were then opened on Imaris 10.1 for visualization and thresholding before defining regions of interest (ROI) for each sample. Twelve brain regions were selected for analysis. Region dimensions were created using the Allen Brain Atlas 3D adult mouse brain viewer (atlas.brain-map.org). This 3D viewer included an adjustable scale bar and predefined region boundaries to enable ROI creation (See Supplementary Data 1 for ROI dimensions). After ROI dimensions were determined, ROIs were drawn on each sample. Then, Imaris spot detection and three nearest neighbor analyses were performed.

Briefly, the spot detection analysis identifies and counts all CX3CR1^GFP+^ cells in each region. After each cell has been identified, the three nearest neighbor analysis measures the distance between each cell and its three nearest neighbor cells. The program then calculates the average distance for all cells and their three nearest neighbors in a given region. The three nearest neighbor analysis enables researchers to identify patterns of cell clustering within and between regions. Further, evidence suggests that this analysis can be used as an indirect measure of microglia and brain macrophage reactivity (Davis et al., 2017). Spot detection and three nearest neighbor analyses were performed in ROIs of both hemispheres. CX3CR1^GFP+^ cell count and three nearest neighbor data for both hemispheres were averaged within each animal.

Due to the varying sizes of the ROIs, cell counts for each region were normalized to the number of cells per mm^3^. Data was then exported and uploaded to Graphpad Prism 10 for all statistical analyses.

### 2.9 Statistical Analyses

Mixed-effects analyses were utilized to measure sex and exposure-dependent changes in binge-like ethanol consumption (Figure 2; Supplementary Data 2: Figures 1-3). Two-way ANOVA analyses were utilized to identify region and exposure-dependent changes in cell count and three nearest neighbor proximity between regions; and exposure and sex-dependent changes in cell count and three nearest neighbor proximity within regions (Figure 3 f, i; Figure 4; Supplementary Data 2: Figures 5-8).

**Figure 2.**
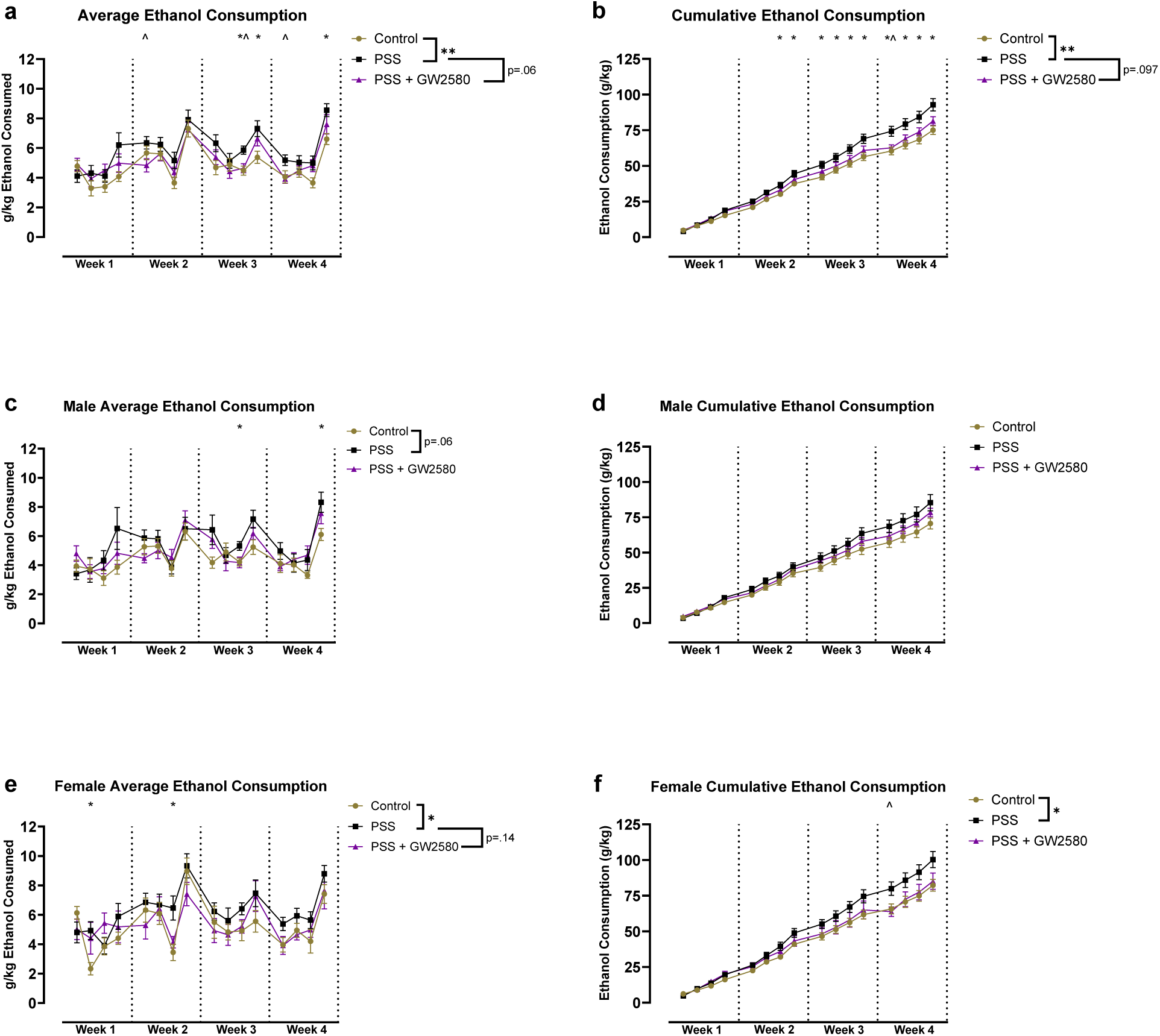
PSS increases binge-like ethanol consumption. a) Average daily ethanol consumption for control, PSS-exposed, and GW2580-treated mice. b) Cumulative ethanol consumption for control, PSS-exposed, and GW2580-treated mice. c) Average daily ethanol consumption for male mice. d) Cumulative ethanol consumption for male mice. e) Average daily ethanol consumption for female mice. f) Cumulative ethanol consumption for female mice. Error bars indicate mean ± s.e.m. * Indicates days that PSS-exposed mice consumed more ethanol than control mice. ^ indicates days that PSS-exposed mice consumed more ethanol than GW2580-treated mice. * Indicates p<.05; ** indicates p<.01; *** indicates p<.001; and **** indicates p<.0001.

**Figure 3.**
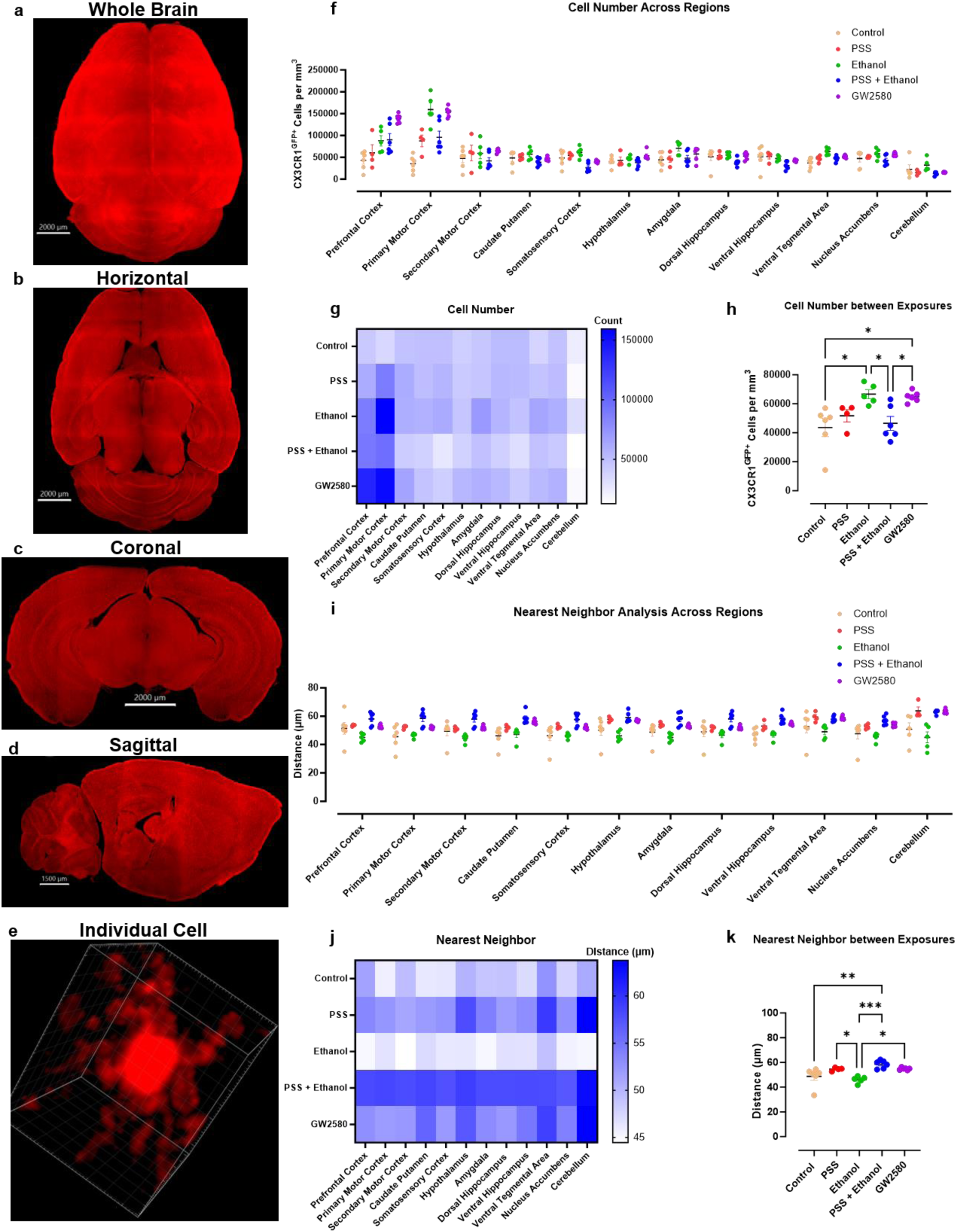
GW2580 treatment prevents PSS + ethanol-induced changes in microglia and brain macrophage count. a-e) Representative images of a 3D reconstructed brain and an individual CX3CR1^GFP+^ cell. f) Average CX3CR1^GFP+^ cell count for each exposure within each region of the brain. g) Heat map providing region x exposure comparisons for CX3CR1^GFP+^ cell count. h) The average CX3CR1^GFP+^ cell count for each brain within each exposure condition. i) Average CX3CR1^GFP+^ cell proximity for each exposure within each region of the brain. j) Heat map providing region x exposure comparisons for CX3CR1^GFP+^ cell proximity. k) Average CX3CR1^GFP+^ cell proximity for each brain within each exposure condition. Error bars indicate mean ± s.e.m. * Indicates p<.05; ** indicates p<.01; *** indicates p<.001; and **** indicates p<.0001.

**Figure 4.**
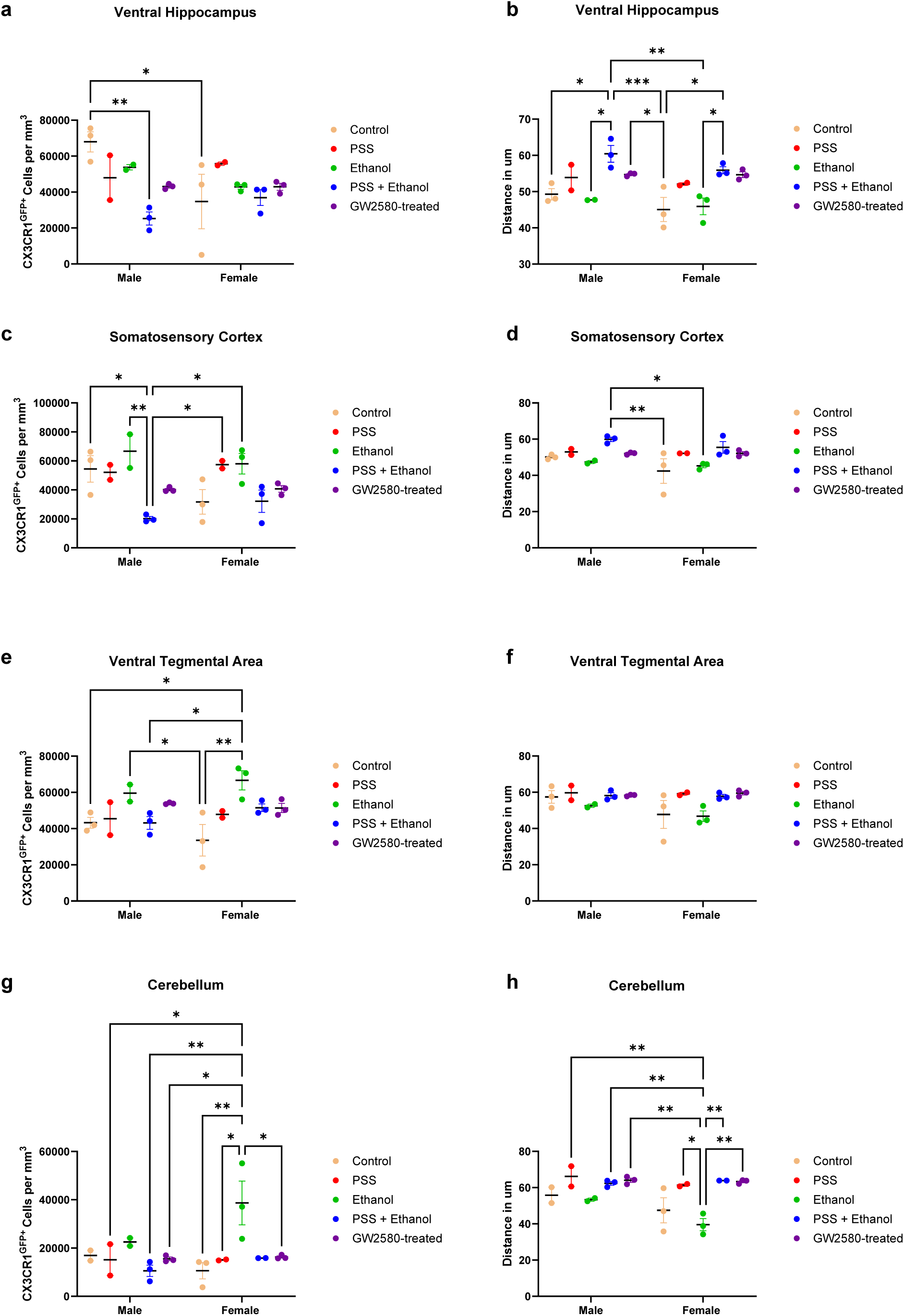
PSS and repeated binge-like ethanol consumption alter microglia and brain macrophage count and proximity in a sex and region-dependent manner. a) CX3CR1^GFP+^ cell count in the ventral hippocampus. b) CX3CR1^GFP+^ cell proximity in the ventral hippocampus. c) CX3CR1^GFP+^ cell count in the somatosensory cortex. d) CX3CR1^GFP+^ cell proximity in the somatosensory cortex. e) CX3CR1^GFP+^ cell count in the ventral tegmental area. f) CX3CR1^GFP+^ cell proximity in the ventral tegmental area. g) CX3CR1^GFP+^ cell count in the cerebellum. h) CX3CR1^GFP+^ cell proximity in the cerebellum. Error bars indicate mean ± s.e.m. * Indicates p<.05; ** indicates p<.01; *** indicates p<.001; and **** indicates p<.0001.

One-way ANOVA analyses were performed to determine the effect of exposure or region on cell count and proximity (Figure 3h, k; Supplementary Data 2: Figure 4). Tukey’s post-hoc comparisons were performed when main effects or interactions were identified following any of the above mentioned analyses. Correlations were performed to determine the strength of the relationship between CX3CR1^GFP+^ count and proximity (Supplementary Data 2: Figures 9-10). All statistics were performed using Graphpad Prism 10. If interested in a set of statistical outcomes, see Supplementary Data 3.

## 3. Results

### 3.1 Early life PSS increases binge-like ethanol consumption

The primary goal of this study was to determine whether early life PSS would alter binge-like ethanol consumption in adulthood. Further, we sought to identify whether pharmacological inhibition of CSF1Rs on microglia and brain macrophages during PSS would alter or prevent the effect of PSS on binge-like ethanol consumption. Mice were assigned to either control + vehicle (n=21; 13 male and 8 female), PSS + vehicle (n=18; 9 male and 9 female), or PSS + GW2580 (n=18; 10 male and 8 female). At P60, CX3XR1^GFP+^ mice began DID (Figure 1).

A mixed-effects analysis revealed main effects of exposure (p=.006) and time (p<.0001) on average daily ethanol consumption, but no exposure x time interaction (p=.19). PSS-exposed mice consumed more ethanol than control mice (p=.003). Further, there was a strong trend towards PSS-exposed mice drinking more ethanol than GW2580-treated mice (p=.06) (Figure 2a). We then identified sex-dependent effects of PSS on binge-like ethanol consumption. In male mice, a mixed-effects analysis demonstrated a main effect of time (p<.0001), a trend towards a main effect of exposure (p=.09), but no exposure x time interaction (p=.15). However, there was a trend towards PSS-exposed male mice consuming more ethanol than control male mice (p=.06) (Figure 2c). Consistent with male mice, a mixed-effects analysis revealed a main effect of time (p<.0001), a trend towards a main effect of exposure (p=.08), but no exposure x time interaction (p=.32) on average daily ethanol consumption in female mice. However, PSS-exposed female mice consumed more ethanol than control mice (p=.029), and PSS-exposed female mice trended towards having higher average daily ethanol consumption than GW2580-treated female mice (p=.14) (Figure 2e).

We next sought to determine the effect of these exposures on cumulative ethanol consumption. A mixed-effects analysis revealed main effects of exposure (p=.015) and time (p<.0001); and an exposure x time interaction (p<.0001). PSS-exposed mice consumed more ethanol than control mice (p=.0064), and trended towards consuming more ethanol than GW2580-treated mice (p=.097) (Fig 2b). In male mice, a mixed-effects analysis demonstrated a main effect of time (p<.0001), no effect of exposure (p=.17), and an exposure x time interaction (p<.0001) (Figure 2d). In female mice, a mixed-effects analysis revealed a main effect of time (p<.0001), a trend towards an effect of exposure (p=.12), and an exposure x time interaction (p<.0001). However, PSS-exposed female mice consumed more ethanol across the study when compared to control female mice (p=.036) (Figure 2f). Overall, we found that PSS increased binge-like ethanol consumption. GW2580 treatment trended towards blunting the effect of PSS on binge-like ethanol consumption. Further, PSS and GW2580 treatment had a more robust effect on female mice when compared to male mice. For all statistics and additional group comparisons view Supplementary Data 2: Figures 1-3 and Supplementary Data 3.

### 3.2 GW2580 treatment prevents the effects of PSS on microglia and brain macrophage count following repeated binge-like ethanol consumption

Following completion of the DID procedure, brains (n=27) were dissected from control (n=6; 3 male and 3 female), PSS-exposed (n=4; 2 male and 2 female), ethanol-exposed (n=5; 2 male and 3 female), PSS + ethanol-exposed (n=6; 3 male and 3 female), and GW2580-treated (n=6; 3 male and 3 female) mice for LSFM (Figure 3a-e; Supplementary Data 4-8). We sought to determine how control, PSS, repeated binge-like ethanol consumption, the combination of PSS and repeated binge-like ethanol consumption, and GW2580 treatment during PSS altered microglia and brain macrophage population densities across twelve brain regions including the prefrontal cortex, primary motor cortex, secondary motor cortex, somatosensory cortex, caudate putamen, hypothalamus, nucleus accumbens, amygdala, dorsal hippocampus, ventral hippocampus, ventral tegmental area, and cerebellum. Regions were selected to ensure adequate coverage of the brain; and based on their relevancy to stress-responsivity, reward learning, and motivated behaviors. First, Imaris spot detection analysis was performed to count the number of CX3CR1^GFP+^ microglia and brain macrophages within each region. Second, a nearest neighbor analysis was performed to identify the average distance between each cell and its three nearest neighbors.

When investigating differences in microglia and brain macrophage count between regions, a two-way ANOVA revealed main effects of exposure (p<.0001) and region (p<.0001) as well as an exposure x region interaction (p<.0001). Post-hoc analyses demonstrated that microglia and brain macrophage count were highest in the prefrontal cortex and primary motor cortex. In addition, these regions were much more responsive to each exposure when compared to other investigated regions. Consistent with prior work (Savchenko et al., 1997), the cerebellum exhibited significantly lower microglia and brain macrophage count (Figure 3f-g; Supplementary Data 2: Figure 4; Supplementary Data 3).

Next, we averaged the number of microglia and brain macrophages observed in each region for each brain and compared the effect of each exposure on cell count. A one-way ANOVA revealed a significant effect of exposure on microglia and brain macrophage count (p=.0023). Post-hoc analyses demonstrated that control mice had less microglia and brain macrophages than ethanol-treated (p=.01) and GW2580-treated (p=.01) mice. Further, PSS + ethanol-exposed mice had less microglia and brain macrophages when compared to ethanol-treated (p=.03) and GW2580-treated (p=.04) mice (Figure 3h). This suggests that GW2580 treatment prevented PSS-induced changes in cell count following repeated binge-like ethanol consumption.

We then assessed whether microglia and brain macrophage proximities were exposure and region-dependent. A two-way ANOVA revealed main effects of exposure (<.0001) and region (p<.0001), but no interaction (Figure 3i-j, Supplementary Data 2: Figure 4). Following completion of this analysis, we averaged the nearest neighbor data collected from each region for each brain. A one-way ANOVA revealed that the exposures had an effect on cell proximity (p=.0004). Post-hoc analyses demonstrated that control mice had less distance between cells than PSS + ethanol-exposed mice(.005). Further, ethanol-treated mice had less distance between cells than PSS-exposed (p=.05), PSS + ethanol-exposed (p=.0006), and GW2580-treated (p=.02) mice (Figure 3k). Our data suggest that PSS altered microglia and brain macrophage proximity following repeated binge-like ethanol consumption when compared to repeated binge-like ethanol consumption alone. GW2580 treatment did not prevent this outcome from occurring.

### 3.3 PSS and repeated binge-like ethanol consumption alter CX3CR1^GFP+^ microglia and brain macrophage population density in a sex and region-dependent manner

Following completion of analyses identifying region and exposure-dependent changes in CX3CR1^GFP+^ cell count and proximity across and between brain regions, we turned our attention to investigating exposure and sex-dependent changes in CX3CR1^GFP+^ cell count and proximity within each region. Although we found interesting exposure and sex-dependent effects on count and proximity in most regions (Supplementary Data 2: Figures 5-8), we focused our commentary on the ventral hippocampus, somatosensory cortex, ventral tegmental area, and cerebellum.

In the ventral hippocampus, a two-way ANOVA revealed a main effect of exposure (p=.04), no effect of sex (p=.27), and an exposure x sex interaction (p=.02). Post-hoc analyses demonstrated control male mice had more CX3CR1^GFP+^ cells than PSS + ethanol-exposed male (p=.006) and control female (p=.05) mice (Figure 4a). Further, a two-way ANOVA revealed a main effect of exposure (p<.0001), a trend towards a main effect of sex (p=.07), and no exposure x sex interaction (p=.75) on microglia and brain macrophage proximity. PSS + ethanol-exposed male mice had greater distance between CX3CR1^GFP+^ cells than control (male: p=.02; female: p=.0007) and ethanol-treated (male: p=.01; female: p=.001) mice. GW2580-treated male mice had greater distance between CX3CR1^GFP+^ cells than control female mice (p=.05).

While PSS + ethanol-exposed female mice exhibited greater distance between CX3CR1^GFP+^ cells than control (p=.02) and ethanol-treated (p=.04) female mice (Figure 4b). Our data suggest PSS reduced microglia and brain macrophage count following repeated binge-like ethanol consumption and GW2580 treatment prevented the effect of PSS on microglia and brain macrophage count in male mice. Further, GW2580 treatment prevented PSS-evoked changes in microglia and brain macrophage proximity following repeated binge-like ethanol consumption

In the somatosensory cortex, a two-way ANOVA revealed a main effect of exposure (p=.0005), no effect of sex (p=.52), and a trend towards an exposure x sex interaction (p=.11) on CX3CR1^GFP+^ cell count. Post-hoc analyses demonstrated that PSS + ethanol-exposed male mice had fewer CX3CR1^GFP+^ cells than control (p=.03) and ethanol-treated (p=.006) male mice as well as PSS-exposed (p=.04) and ethanol-treated (p=.01) female mice (Figure 4c). Further, a two-way ANOVA revealed a main effect of exposure (p=.003), but not sex (p=.12) or an exposure x sex interaction (p=.64) on microglia and brain macrophage proximity. Post-hoc analyses demonstrated that PSS + ethanol-exposed male mice had greater distance between CX3CR1^GFP+^ cells than control (p=.007) and ethanol-treated (p=.03) female mice (Figure 4d). Our data suggest PSS reduced microglia and brain macrophage count following repeated binge-like ethanol consumption and GW2580 treatment prevented the effect of PSS on microglia and brain macrophage count in male mice.

In the ventral tegmental area, a two-way ANOVA revealed a main effect of exposure (p=.001), but not sex (p=.72) or an exposure x sex interaction (p=.31) on CX3CR1^GFP+^ cell count. Post-hoc analyses demonstrated ethanol-treated male mice had more CX3CR1^GFP+^ cells than control female mice (p=.05). While ethanol-treated female mice had more CX3CR1^GFP+^ cells than control (male: p=.04; female: p=.002) mice and PSS + ethanol-exposed (p=.04) male mice (Figure 4e). Further, a two-way ANOVA revealed a main effect of exposure (p=.05), but not sex (p=.19) or an exposure x sex interaction (p=.47) on microglia and brain macrophage proximity. Post-hoc analyses demonstrated no further group differences (Figure 4f).

In the cerebellum, a two-way ANOVA revealed main effects of exposure (p=.006), but not sex (p=.27) or an exposure x sex interaction (p=.18) on CX3CR1^GFP+^ cell count. Post-hoc analyses demonstrated ethanol-treated female mice had more CX3CR1^GFP+^ cells than PSS-exposed (p=.04), PSS + ethanol-exposed (p=.004), and GW2580-treated (p=.02) male mice as well as control (p=.004), PSS-exposed (p=.04), and GW2580-treated (p=.03) female mice (Figure 4g). Further, a two-way ANOVA revealed main effects of exposure (p=.0003) and sex (p=.04), but no exposure x sex interaction (p=.26) on microglia and brain macrophage proximity. Post-hoc analyses demonstrated that ethanol-treated female mice had shorter distance between CX3CR1^GFP+^ cells than PSS-exposed (male: p=.003; female: p=.02), PSS + ethanol-exposed (male: p=.005; female: p=.007), and GW2580-treated (male: p=.002; female: p=.003) mice (Figure 4h).

## 4. Discussion

In this study, we sought to determine the effect of an ELS model, PSS, on binge-like ethanol consumption; identify changes in microglia and brain macrophage population densities in twelve brain regions; and determine whether microglia and brain macrophage inhibition could prevent PSS-evoked changes in binge-like ethanol consumption and alterations in microglia and brain macrophage population densities.

We found that PSS increased binge-like ethanol consumption in early adulthood, while inhibiting microglia and brain macrophage proliferation during PSS trended towards reducing PSS-evoked increases in binge-like ethanol consumption. After looking at the sexes independently, we found that PSS had a more robust effect on female mice and GW2580 treatment also appeared to be more effective at blunting the effects of PSS on binge-like ethanol consumption in female mice. A growing body of evidence suggests that various forms of ELS can facilitate sex-dependent outcomes on behavior (Bath, 2020; Catale et al., 2020). While previous work utilizing PSS (Lo Iacono et al., 2016, 2017, 2018; Catale et al., 2020, 2022), identified changes in behavior, brain vasculature, dopamine neuron excitability, and microglia, none of their work explored sex-differences. Our data suggest that PSS induces sex-dependent changes in binge-like ethanol consumption, and microglia and brain macrophage population densities. As such, future work should consider biological sex as a variable when utilizing PSS.

Recent efforts in the alcohol research community have advanced pharmacotherapies targeting inflammation to treat AUD (Grodin et al., 2021). While preclinical evidence suggests that targeting microglia can prevent alcohol dependence (Warden et al., 2020), no study has demonstrated that targeting microglia can treat AUD. With this in mind, perhaps it is time we consider taking a preventative approach to AUD. Our study utilized GW2580, a CSF1R inhibitor that prevents microglia and brain macrophage proliferation (Neal et al., 2020). While acute CSF1R inhibition has a direct effect on proliferation, it has not been directly linked to microglial and brain macrophage engagement with synapses and neurons. We found that CSF1R inhibition trended towards reducing binge-like ethanol consumption following PSS. We speculate that this approach to inhibiting microglia and brain macrophages was too generalized and developing a target to modulate microglia-neuron interactions may be a better approach to prevent the effects of PSS on binge-like ethanol consumption.

Following completion of DID, we created 3D reconstructions of CX3CR1^GFP+^ brains from each exposure to identify changes in microglia and brain macrophage population density across the brain. We found that repeated binge-like ethanol consumption, but not PSS, increased the number of CX3CR1^GFP+^ cells. However, early life PSS blunted ethanol-induced changes in CX3CR1^GFP+^ cell count. Our data suggest that early life PSS altered microglia and brain macrophage responsivity to repeated binge-like ethanol consumption. Previous work has demonstrated that microglia and brain macrophages develop an innate immune memory following a stress exposure resulting in altered function in response to future environmental stimuli (Frank et al., 2014; Neher & Cunningham, 2019, Reemst et al., 2022). Further, recent work has speculated that this process, colloquially referred to as microglial priming, may play a role in the development of AUD (Melbourne et al., 2021).

Our data offer some support for this hypothesis. GW2580 was sufficient to prevent the effects of PSS on microglia and brain macrophage count following repeated binge-like ethanol consumption. While this finding was exciting, GW2580 treatment did not prevent the effect of PSS on microglia and brain macrophage proximity. As such, our data suggest that CSF1Rs may play a role in innate immune memory formation, but does not facilitate the entirety of the process. Although we were excited to find this outcome when looking at these brain regions collectively, microglia and brain macrophages are present heterogeneously throughout the brain (Savchenko et al., 1997; Tan et al., 2019). Therefore, we were interested in identifying how individual regions responded to each exposure. While we found interesting outcomes in most regions, we decided to discuss four regions in particular: the ventral hippocampus, the somatosensory cortex, the ventral tegmental area, and the cerebellum.

First, we identified how biological sex and these environmental exposures interacted to alter microglia and brain macrophage count and proximity in the ventral hippocampus. The ventral hippocampus acts as a major relay point between numerous regions involved with emotion regulation, motivation, and reward (Turner et al., 2022). ELS has been demonstrated to have lasting consequences on ventral hippocampal plasticity (Çalışkan et al., 2020). Further, chronic intermittent ethanol exposure alters neuronal excitability in the ventral hippocampus resulting in increased anxiety-like behaviors (Ewin et al., 2019, Bach et al., 2021). Due to its role in regulating behaviors related to ELS and AUD, we determined the ventral hippocampus to be an ideal candidate to include in our discussion.

We found that control male mice had higher CX3CR1^GFP+^ cell count when compared to PSS + ethanol-exposed male mice and control female mice. Previous work has demonstrated that repeated binge-like ethanol consumption reduces microglia and brain macrophage count in the dorsal and ventral hippocampus (Nelson et al., 2021).

While we did not replicate these findings, we propose the difference in outcomes is, in part, due to the difference in scale between the observations. LSFM can provide whole region readouts, and may be more comprehensive than traditional microscopy approaches that do not consider the region in its entirety. In spite of this failure to replicate previous findings, our data demonstrate that GW2580 prevented the effect of PSS on microglia and brain macrophage count following repeated binge-like ethanol consumption in male mice. We then performed a three nearest neighbor analysis in the ventral hippocampus. Regardless of sex, PSS + ethanol-exposed mice exhibited greater distance between cells when compared to ethanol alone mice. However, GW2580 treatment prevented the effect of PSS on microglia and brain macrophage count and proximity following repeated binge-like ethanol consumption. Data collected from the ventral hippocampus supports the notion that ELS experiences such as PSS may prime microglia to be more reactive to future environmental insults, like ethanol consumption. Further, targeted inhibition of these cells during ELS may prevent long-term adaptations in function.

We then proceeded to perform analyses on the somatosensory cortex. We chose the somatosensory cortex for several reasons. First, ELS is known to alter the developmental trajectory of the somatosensory cortex (Takatsuru & Koibuchi, 2015).

The somatosensory cortex’s primary function is to process sensory information including nociception (Kropf et al., 2018). The PSS exposure we utilized in this study involves a lot of tactile stimulation. Previous work suggests that tactile stimulation between P14-16 can alter longitudinal neuronal activity within the somatosensory cortex (He et al., 2017). Further, alcohol intoxication increases functional connectivity between the somatosensory cortex and the brain stem which is related to issues with coordinating motor function (Shokri-Kojori et al., 2017). If PSS can alter the developmental trajectory of the somatosensory cortex leading to individuals being more prone to injury due to issues with motor impairment while under the influence of alcohol, preventing this outcome via microglia and brain macrophage inhibition during PSS could reduce this risk. We found that male mice that were exposed to PSS and that engaged in repeated binge-like ethanol consumption had fewer CX3CR1^GFP+^ cells than control and ethanol-treated male mice and PSS or ethanol-treated female mice. Mice treated with GW2580 during PSS did not exhibit this change in CX3CR1^GFP+^ cell count suggesting that microglia inhibition was sufficient to prevent this long-term outcome.

We then assessed how PSS and repeated binge-like ethanol consumption altered microglia and brain macrophage population density in the ventral tegmental area. The ventral tegmental area is a key regulator of environmental stimuli salience, reward processing, and the development of AUD (Koob & Volkow, 2010; You et al., 2018). Existing literature widely supports the notion that ELS can blunt ventral tegmental area development leading to issues with reward appraisal (Park et al., 2021). Previous work has also demonstrated that PSS alone does not increase microglia and brain macrophage count in adulthood. However, PSS followed by cocaine administration can increase the number of microglia and brain macrophages within the ventral tegmental area (Lo Iacono et al., 2018).

As a result, the logic behind our decision to investigate the ventral tegmental area was two-fold: 1) this region is susceptible to ELS and is an important regulator of alcohol use and 2) we wanted to determine if we could replicate previous findings. Consistent with prior work, we found that PSS alone did not have an effect on CX3CR1^GFP+^ cell count or proximity in the ventral tegmental area. However, ethanol-treated female mice exhibited an increase in the number of CX3CR1^GFP+^ cells when compared to male and female control mice and PSS and ethanol-treated male mice. Interestingly, the combination of PSS and ethanol consumption did not lead to an increase in ventral tegmental area microglia and brain macrophage count, as observed in PSS mice administered cocaine later in life. This finding is not at surprising as there are profound differences in the pharmacokinetic and pharmacodynamic properties of cocaine and alcohol. Although both cocaine and alcohol can increase inflammation (Sil et al., 2019; van de Loo et al., 2020), it is likely that the mechanisms underlying their pro-inflammatory effects are distinct and thus explain their differential effects on microglia and brain macrophage count observed between the studies.

The last region we included in this discussion is the cerebellum. Although the cerebellum is most widely known for its involvement in motor function, a growing body of work suggests that it also plays central roles in executive function, affective processing, and the establishment of motivational behaviors (Moulton et al., 2014). ELS is known to alter the developmental trajectory of the cerebellum in a sex-dependent manner (Moussa-Tooks et al., 2020). Similar to other regions of the brain, alcohol consumption increases inflammation and promotes cell death within the cerebellum (Mitoma et al., 2021). Due to the relationship between the somatosensory cortex and cerebellum in coordinating motor function and the suggested role of the cerebellum in executive function and reward, we determined this region to be a good addition to our discussion.

We found that ethanol-treated female mice had more CX3CR1^GFP+^ cells and less distance between cells when compared to other exposure conditions. We were surprised to find such a robust increase in the number of CX3CR1^GFP+^ cells exclusively in female mice. Recent work suggests that adolescent intermittent ethanol exposure does not cause sex-dependent changes in cerebellar gene expression (Healey et al., 2023). However, adolescent intermittent ethanol exposure causes long-term motor impairment in female rats due to cerebellar damage (de Oliveira et al., 2024). To our knowledge, this is the first report to demonstrate sex-dependent ethanol-induced changes in microglia and brain macrophages in the cerebellum. This highlights the importance of identifying how biological sex and various environmental exposures interact to shape behavior and bodily function.

## 5. Conclusion

Our study found that early life PSS increased binge-like ethanol consumption. Microglia and brain macrophage inhibition via GW2580, during PSS, trended towards reducing binge-like ethanol consumption. When looking at the brain regions collectively, GW2580 treatment prevented the effect of PSS on microglia and brain macrophage count following repeated binge-like ethanol consumption. However, the combined and individual effects of PSS and repeated binge-like ethanol consumption on microglia and brain macrophage count and proximity were complex and in many cases sex, exposure, and region-dependent. Future work will seek to better characterize the effects of PSS on microglia and neuron interactions and its consequences on maladaptive drinking behaviors.

## Supporting information

Regions of Interest

Statistics

Nucleus Accumbens

Hippocampus Video

Individual Macrophage

## Funding sources

WF-TARC P50 AA26117; R37 AA17531; F31 AA030928

## CRediT authorship contribution statement

Stephen Gironda: Conceptualization, Data Curation, Formal Analysis, Investigation, Methodology, Project Administration, Visualization, Writing-Original Draft, Writing-Review and Editing

Samuel Centanni: Resources, Software, Supervision, Validation

Jeffrey Weiner: Funding Acquisition, Methodology, Resources, Supervision, Writing-Reviewing and Editing

## Acknowledgements

We would like to thank Dr. Shannon Macauley, Dr. Valeria Carola, and Dr. Amanda Sierra for providing their expertise and assistance. We would like to thank Biorender for providing resources for figure development.

## Notes

### Competing Interest Statement

The authors have declared no competing interest.

### Summary of Updates

We have removed and reformatted the manuscript to primarily focus on our second study

