## Supplemental Figures for "Early life psychosocial stress increases binge-like ethanol consumption and CSF1R inhibition prevents stress-induced alterations in microglia and brain macrophage population density"

Stephen Gironda et al.

All statistics are available in Supplementary Data 3

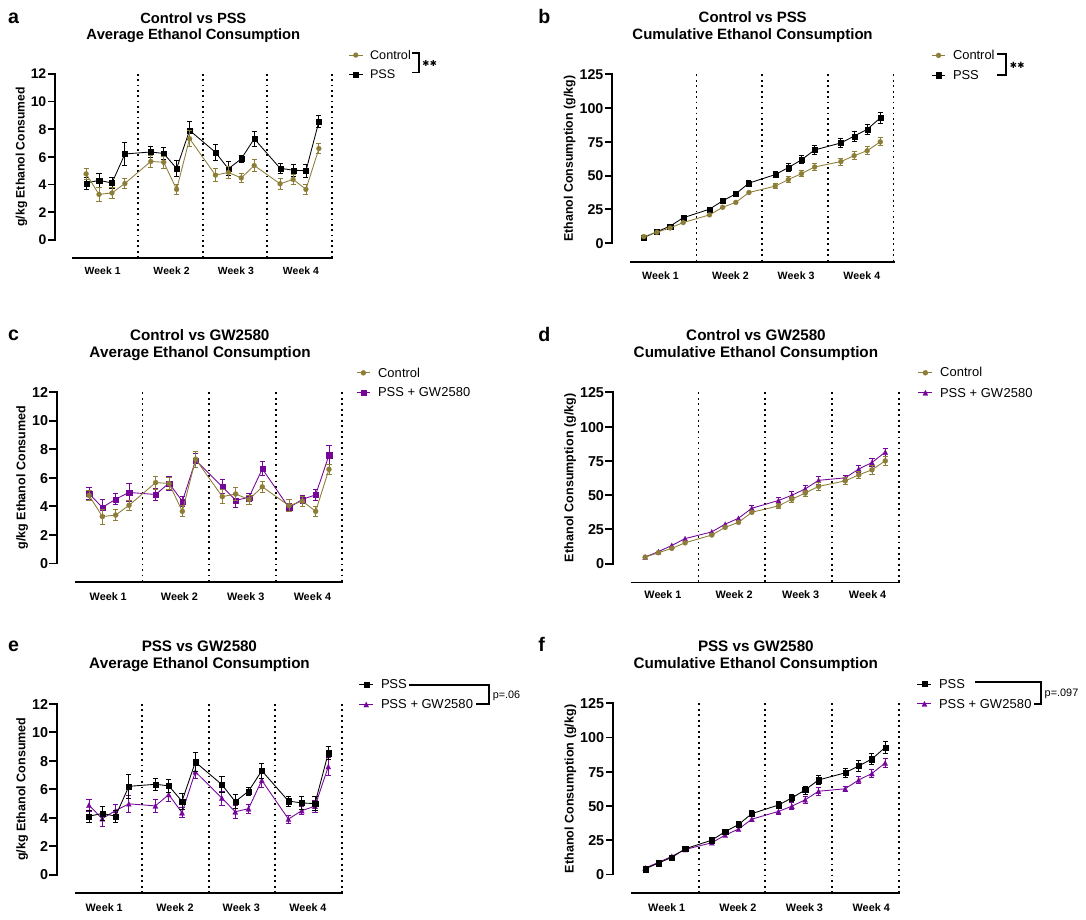

Supplemental Figure 1. PSS-exposed mice consume more ethanol than control CX3CR1^GFP+^ mice. a) Average daily ethanol consumption for control and PSS-exposed mice. b) Cumulative ethanol consumption for control and PSS-exposed mice. c) Average daily ethanol consumption for control and GW2580-treated mice. d) Cumulative ethanol consumption for control and GW2580-treated mice. e) Average daily ethanol consumption for PSS-exposed and GW2580-treated mice. f) Cumulative ethanol consumption for PSS-exposed and GW2580-treated mice. Error bars indicate mean ± s.e.m. * indicates p<.05; ** indicates p<.01; *** indicates p<.001; **** indicates p<.0001.

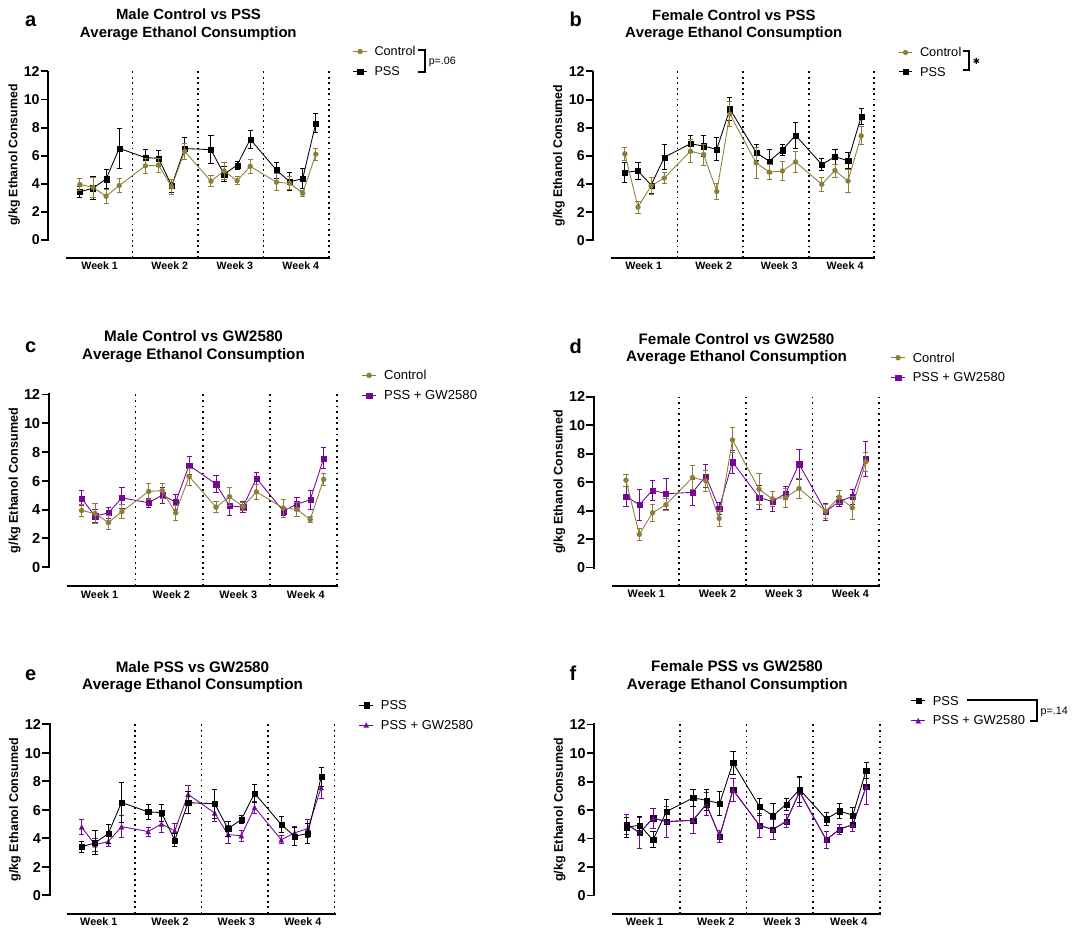

Supplemental Figure 2. Female mice are more vulnerable to the effects of PSS on average daily ethanol consumption. a) Average daily ethanol consumption for male control and PSS-exposed mice. b) Average daily ethanol consumption for female control and PSS-exposed mice. c) Average daily ethanol consumption for male control and GW2580-treated mice. d) Average daily ethanol consumption for female control and GW2580-treated mice. e) Average daily ethanol consumption for male PSS-exposed and GW2580-treated mice. f) Average daily ethanol consumption for female PSS-exposed and GW2580-treated mice. Error bars indicate mean ± s.e.m. * indicates p<.05; ** indicates p<.01; *** indicates p<.001; **** indicates p<.0001.

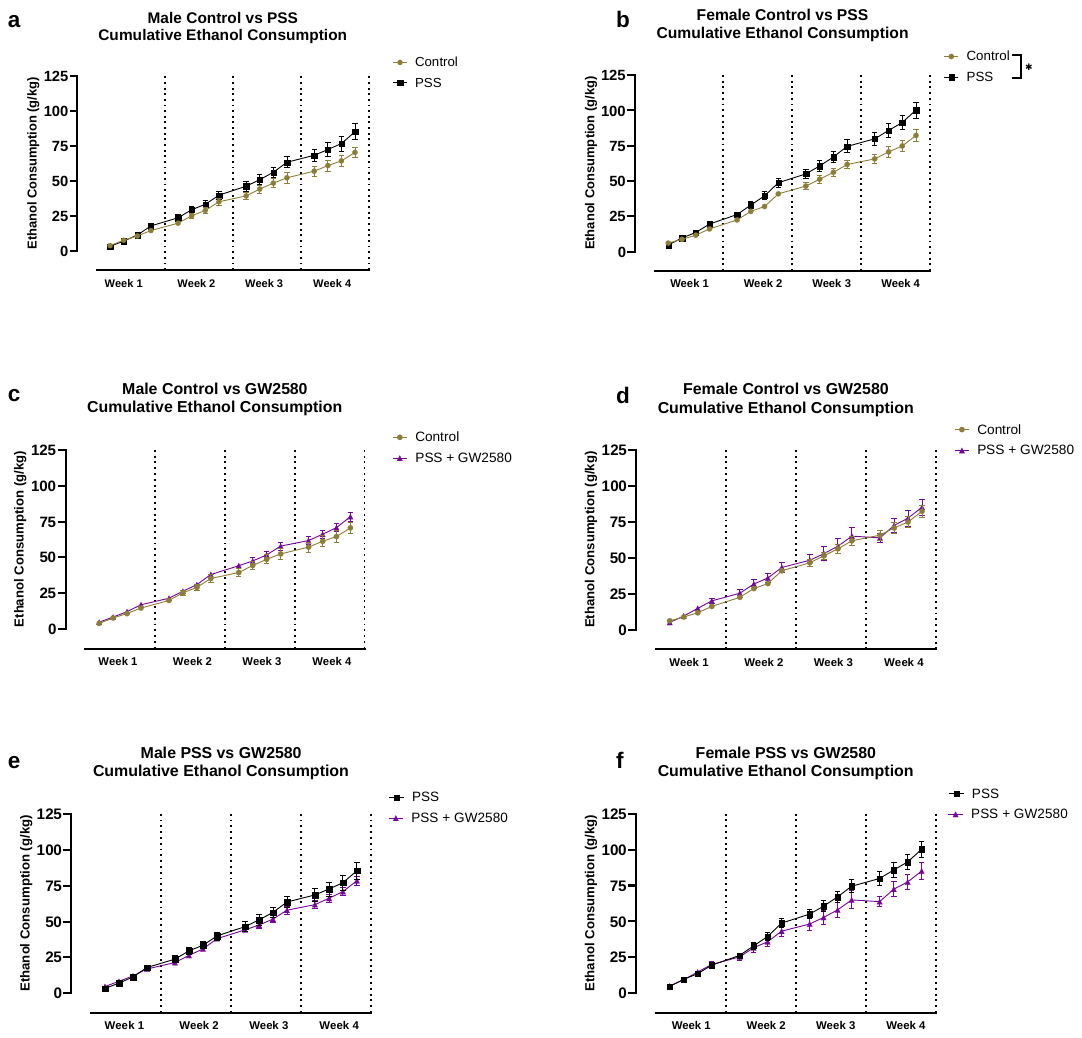

Supplemental Figure 3. Female mice are more vulnerable to the effects of PSS on cumulative ethanol consumption. a) Cumulative ethanol consumption for male control and PSS-exposed mice. b) Cumulative ethanol consumption for female control and PSS-exposed mice. c) Cumulative ethanol consumption for male control and GW2580-treated mice. d) Cumulative ethanol consumption for female control and GW2580-treated mice. e) Cumulative ethanol consumption for male PSS-exposed and GW2580-treated mice. f) Cumulative ethanol consumption for female PSS-exposed and GW2580-treated mice. Error bars indicate mean ± s.e.m. * indicates p<.05; ** indicates p<.01; *** indicates p<.001; **** indicates p<.0001.

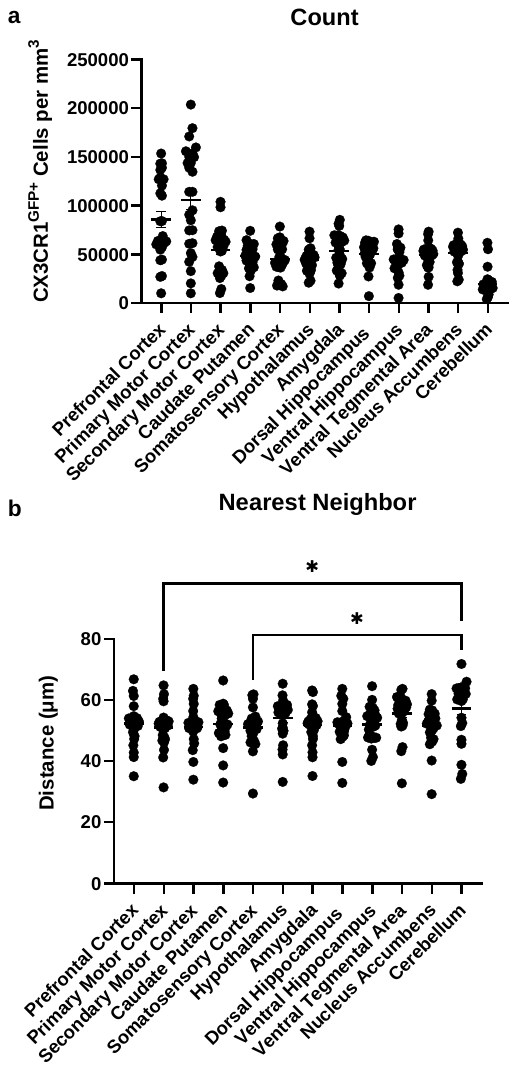

Supplemental Figure 4. Region dependent changes in CX3CR1^GFP+^ cell count and proximity. a) CX3CR1^GFP+^ cell count in each region. b) CX3CR1^GFP+^ cell proximity in each region. There were too many group differences in 4a to plot on the graph (See supplementary data 3 for all pairwise comparisons. Error bars indicate mean ± s.e.m. * indicates p<.05; ** indicates p<.01; *** indicates p<.001; **** indicates p<.0001.

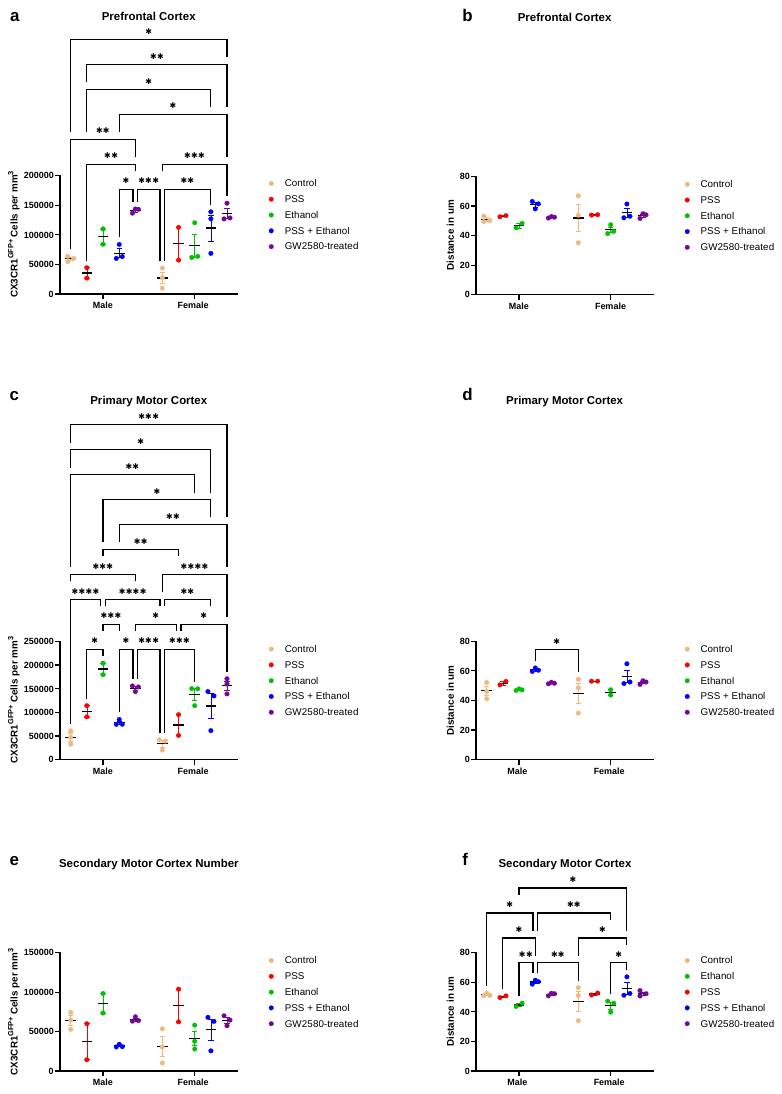

Supplemental Figure 5. CX3CR1^GFP+^ cell count and proximity in the prefrontal cortex, primary motor cortex, and secondary motor cortex. a) The average number of CX3CR1^GFP+^ cells per exposure in the prefrontal cortex. b) The average distance between CX3CR1^GFP+^ cells per exposure in the prefrontal cortex. c) The average number of CX3CR1^GFP+^ cells per exposure in the primary motor cortex. d) The average distance between CX3CR1^GFP+^ cells per exposure in the primary motor cortex. e) The average number of CX3CR1^GFP+^ cells per exposure in the secondary motor cortex. f) The average distance between CX3CR1^GFP+^ cells per exposure in the secondary motor cortex. Error bars indicate mean ± s.e.m. * indicates p<.05; ** indicates p<.01; *** indicates p<.001; **** indicates p<.0001.

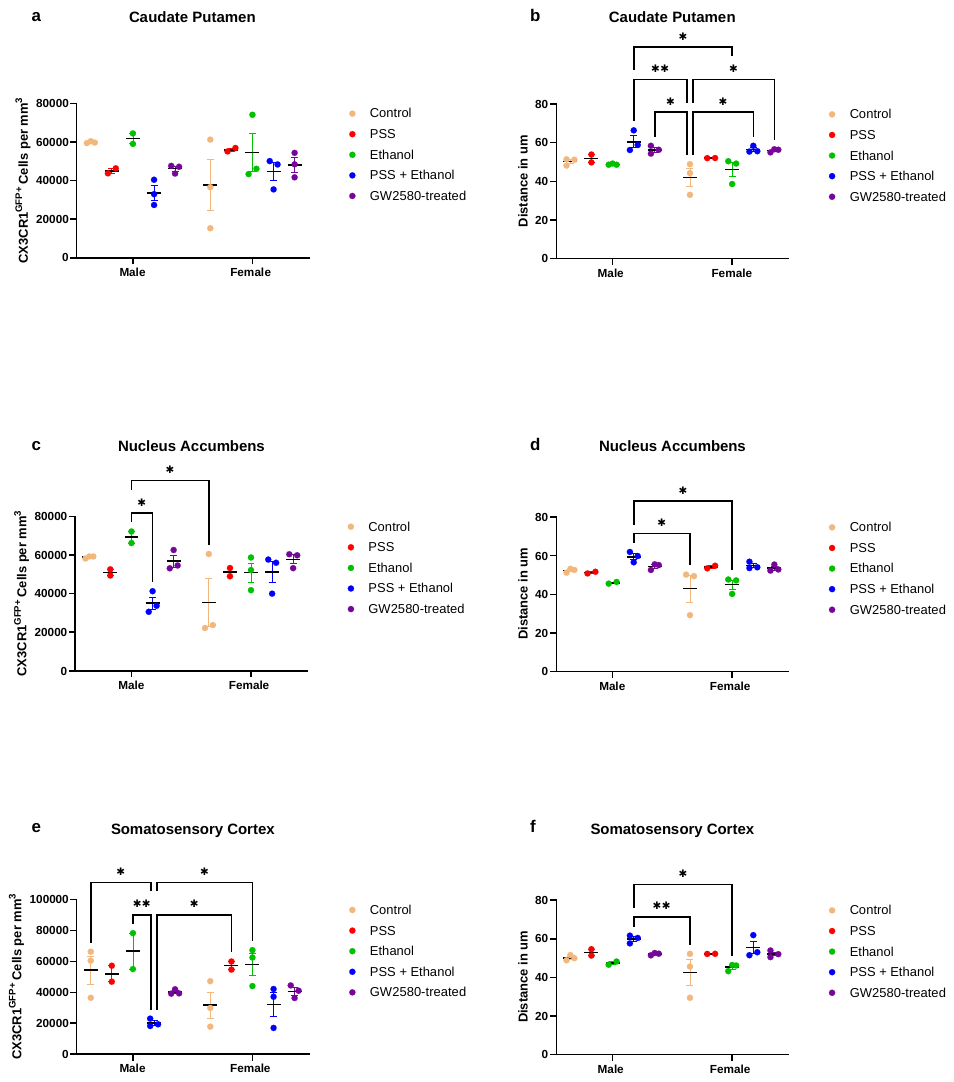

Supplemental Figure 6. CX3CR1^GFP+^ cell count and proximity in the caudate putamen, nucleus accumbens, and somatosensory cortex. a) The average number of CX3CR1^GFP+^ cells per exposure in the caudate putamen. b) The average distance between CX3CR1^GFP+^ cells per exposure in the caudate putamen. c) The average number of CX3CR1^GFP+^ cells per exposure in the nucleus accumbens. d) The average distance between CX3CR1^GFP+^ cells per exposure in the nucleus accumbens. e) The average number of CX3CR1^GFP+^ cells per exposure in the somatosensory cortex. f) The average distance between CX3CR1^GFP+^ cells per exposure in the somatosensory cortex. Error bars indicate mean ± s.e.m. * indicates p<.05; ** indicates p<.01; *** indicates p<.001; **** indicates p<.0001.

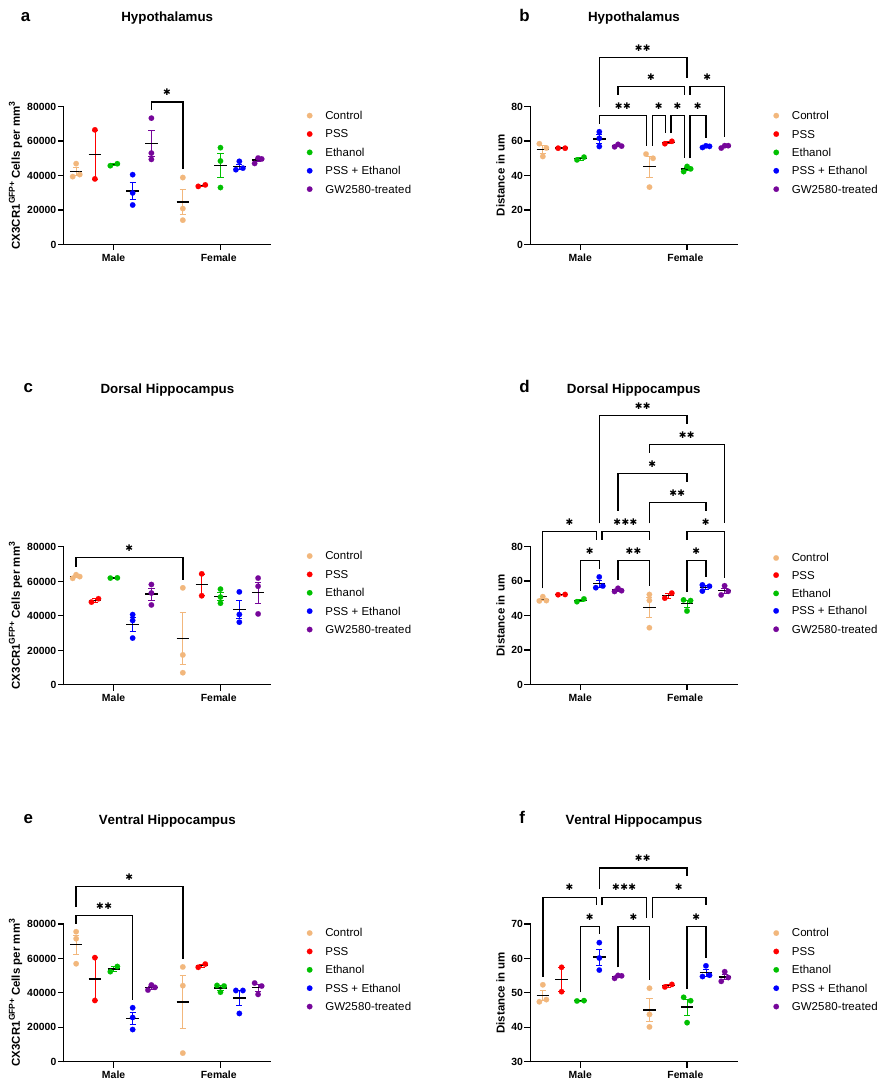

Supplemental Figure 7. CX3CR1^GFP+^ cell count and proximity in the hypothalamus, dorsal hippocampus, and ventral hippocampus. a) The average number of CX3CR1^GFP+^ cells per exposure in the hypothalamus. b) The average distance between CX3CR1^GFP+^ cells per exposure in the hypothalamus. c) The average number of CX3CR1^GFP+^ cells per exposure in the dorsal hippocampus. d) The average distance between CX3CR1^GFP+^ cells per exposure in the dorsal hippocampus. e) The average number of CX3CR1^GFP+^ cells per exposure in the ventral hippocampus. f) The average distance between CX3CR1^GFP+^ cells per exposure in the ventral hippocampus. Error bars indicate mean ± s.e.m. * indicates p<.05; ** indicates p<.01; *** indicates p<.001; **** indicates p<.0001.

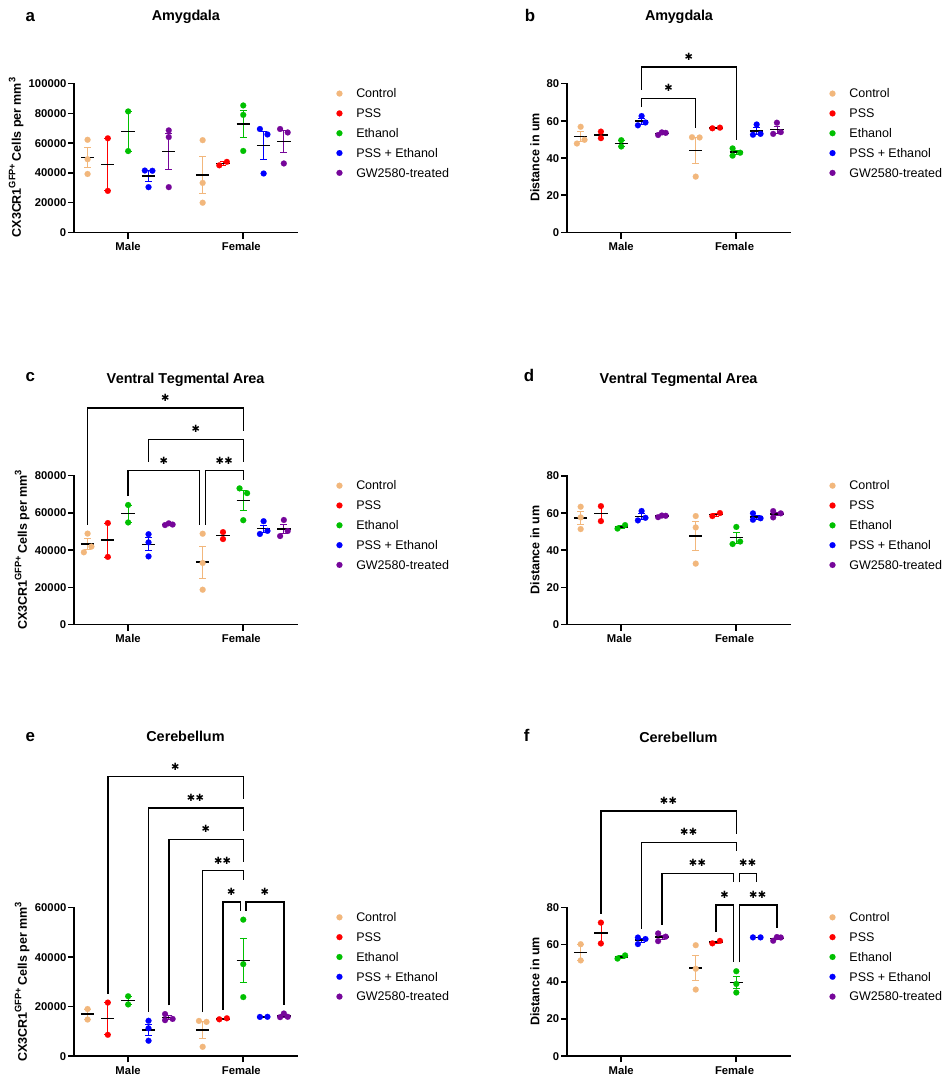

Supplemental Figure 8. CX3CR1^GFP+^ cell count and proximity in the amygdala, ventral tegmental area, and cerebellum. a) The average number of CX3CR1^GFP+^ cells per exposure in the amygdala. b) The average distance between CX3CR1^GFP+^ cells per exposure in the amygdala. c) The average number of CX3CR1^GFP+^ cells per exposure in the ventral tegmental area. d) The average distance between CX3CR1^GFP+^ cells per exposure in the ventral tegmental area. e) The average number of CX3CR1^GFP+^ cells per exposure in the cerebellum. f) The average distance between CX3CR1^GFP+^ cells per exposure in the cerebellum. Error bars indicate mean ± s.e.m. * indicates p<.05; ** indicates p<.01; *** indicates p<.001; **** indicates p<.0001.

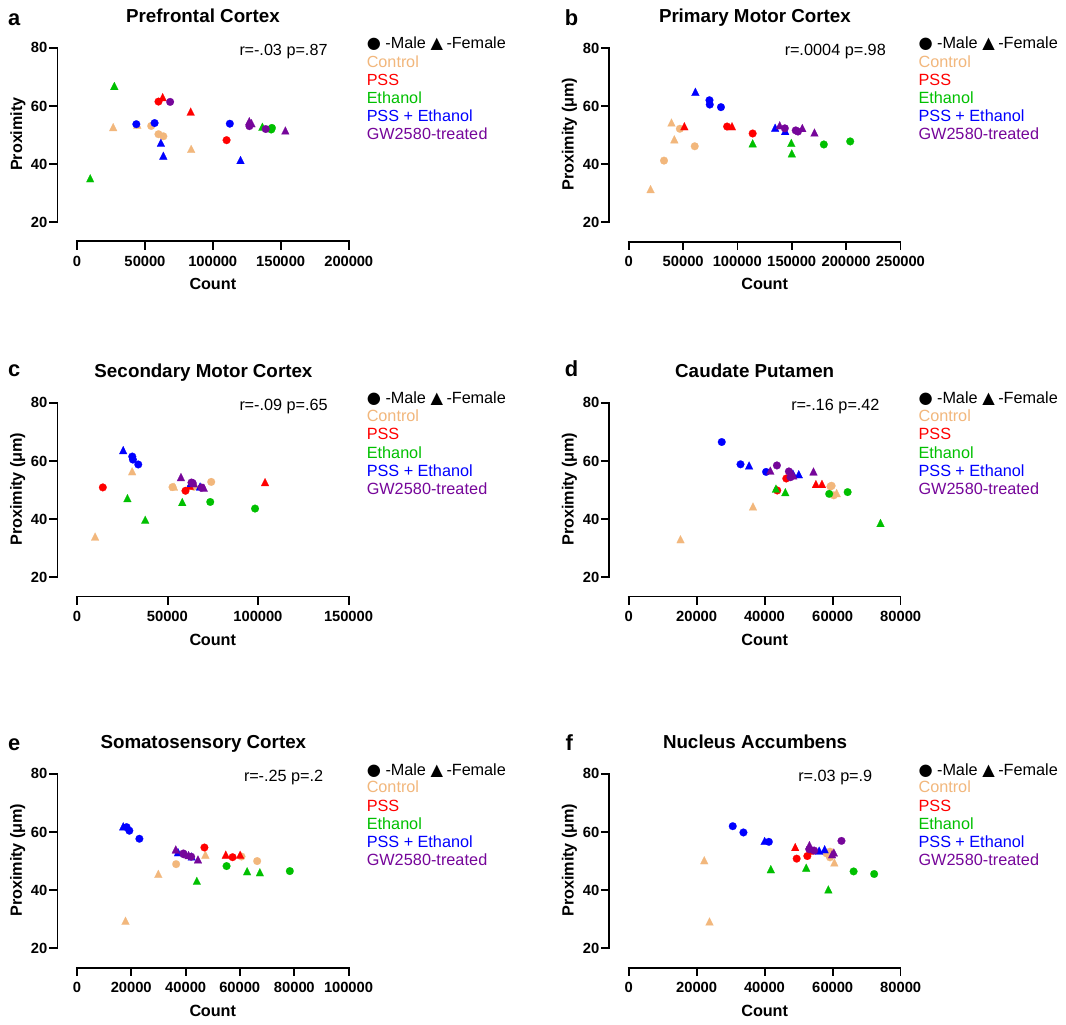

Supplemental Figure 9. Correlations performed to identify the strength of the relationship between count and proximity for each region. a) Relationship between count and proximity in the prefrontal cortex. b) Relationship between count and proximity in the primary motor cortex. c) Relationship between count and proximity in the secondary motor cortex. d) Relationship between count and proximity in the caudate putamen. e) Relationship between count and proximity in the somatosensory cortex. f) Relationship between count and proximity in the nucleus accumbens.* indicates p<.05; ** indicates p<.01; *** indicates p<.001; **** indicates p<.0001.

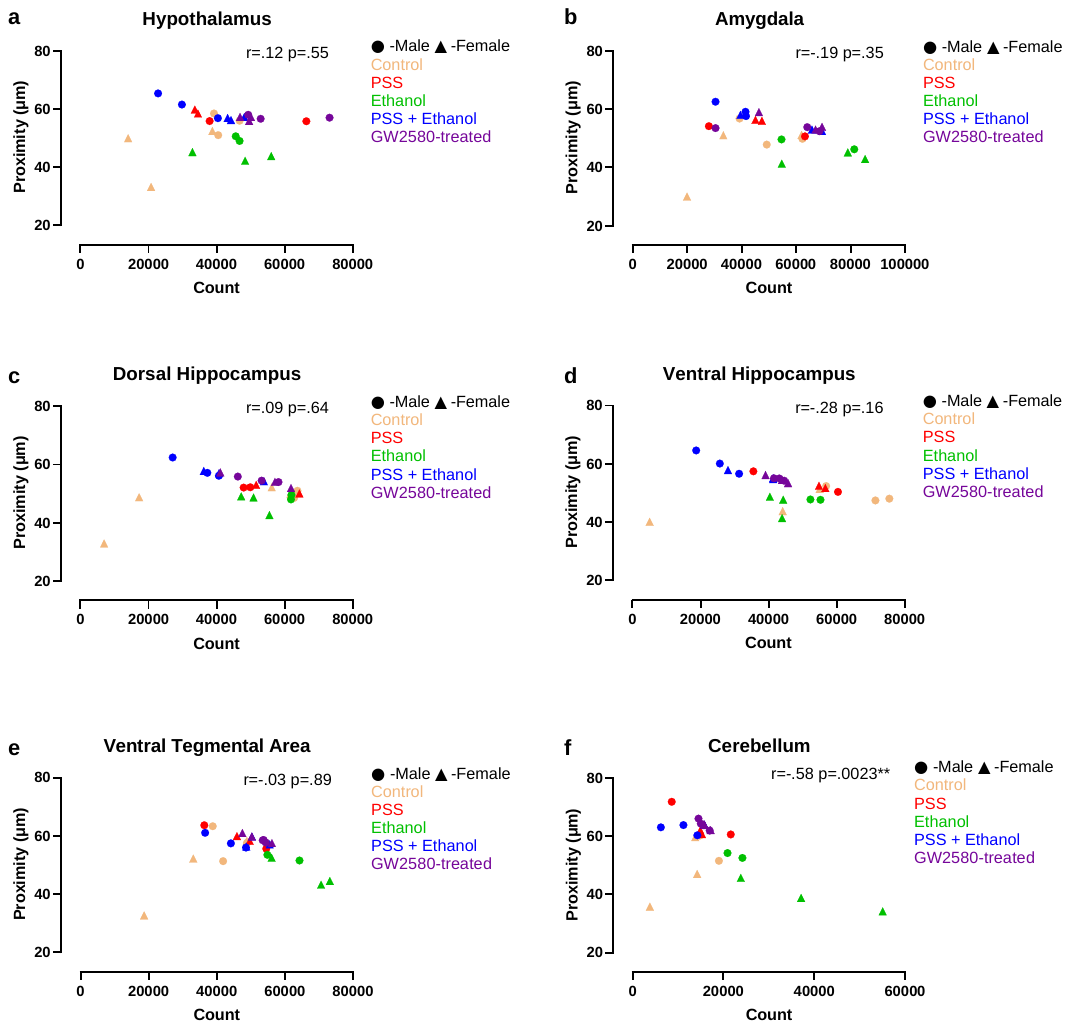

Supplemental Figure 10. Correlations performed to identify the strength of the relationship between count and proximity for each region. a) Relationship between count and proximity in the hypothalamus. b) Relationship between count and proximity in the amygdala. c) Relationship between count and proximity in the dorsal hippocampus. d) Relationship between count and proximity in the ventral hippocampus e) Relationship between count and proximity in the ventral tegmental area. f) Relationship between count and proximity in the cerebellum.* indicates p<.05; ** indicates p<.01; *** indicates p<.001; **** indicates p<.0001.
